# A tumorigenesis threshold for endogenous Myc revealed by dosage-compensation for Myc haploinsufficiency in the absence of p53

**DOI:** 10.1101/2025.07.28.667174

**Authors:** Xiaozhong Bao, Zied Abdullaev, Subhendu K. Das, Valerie Zgonc, Kris Ylaya, Stefania Pittaluga, Stephen M. Hewitt, John M. Sedivy, Svetlana J. Pack, Susan Mackem, David Levens

## Abstract

The *MYC* proto-oncogene is crucial for neoplasia in most tumors. Overexpressed, oncogenic MYC amplifies the flux through most major processes but does not specify a unique carcinogenic pathway. This “amplifier” model suggests that MYC must exceed an expression threshold to become oncogenic. We designed a genetic test of this model, using the mouse *Trp53* null mutant (*p53KO*) as a highly robust tumor generator to examine the effect of a modest change in the endogenous Myc level (*Myc*^+/-^). Strikingly, tumor-free survival is greatly extended in p53KO mice with haploid *Myc* gene-dosage, yet in the tumors that do develop (mainly hemangiosarcomas and thymic lymphomas), their Myc deficit has been invariably compensated either by increasing *Myc* genomic dosage (hemangiosarcomas) or expression (lymphomas). Furthermore, acutely halving the endogenous *Myc* gene-dosage in established tumor allografts curtails growth rates. These results indicate that even an incremental reduction of MYC activity can be salutary in cancer and that one of the major tumor suppressor functions of p53 derives from its ability to prevent *MYC* overexpression. Myc generates acute DNA damage by several mechanisms and accordingly, p53’s anti-Myc function may be inextricably linked to its role in genome integrity surveillance.

## Introduction

The oncogenic actions of MYC (1, 2) and the loss or mutation of tumor suppressor p53 (3-6) contribute to almost all human cancers. Their carcinogenic cooperation has been inferred from the frequency with which they are co-dysregulated in tumors (7-10). Most oncoproteins are activated by mutations that alter their amino-acid sequence to generate qualitatively distinct proteins. In contrast, wild-type MYC protein becomes oncogenic when present in excess (1, 11-13). Increased MYC levels often arise due to gene amplification, chromosomal rearrangements juxtaposing *MYC* with immunoglobulin or other super-enhancers, viral insertions or other transcription-activating signals ectopically transduced to the *MYC* locus, or by mutations that stabilize the normally short-lived MYC protein (14) or RNA (15). Irrespective of the deregulating insult, the result is excess MYC activity. As its levels rise, MYC increasingly binds at all active promoters, amplifying transcription until output saturates (16-19). Once the most active genes become MYC-saturated, excess MYC backfills weaker promoters (17). In this manner, the throughput of all metabolic and signaling pathways are increased in MYC over-producing cells. Thus, MYC may exaggerate whatever programs are triggered by other oncogenes and this oncogenic impulse is abetted by mutations inactivating tumor suppressors. Because of saturation and non-linear transfer functions, the balance between physiological and pathological cellular programs may be disturbed as MYC levels rise (12, 19).

p53 is a master regulator of the cell’s response to genotoxic stress. After DNA damage, p53 may arrest the cell cycle to enable genome repair, or with more extensive damage, may trigger cellular senescence or apoptosis (3-5, 20). In the absence of p53, cells replicate through non-cell lethal damage, progressively accumulating further mutations and chromosomal aberrations that drive tumorigenesis. Mice lacking *Trp53* in the germline rapidly and inevitably develop cancers, mainly lymphomas, hemangiosarcomas, osteosarcomas, and germ-cell tumors (depending partly on strain) (21). Organ-specific *Trp53*-loss provokes tumorigenesis within the targeted tissue, especially when paired with enforced expression of oncogenes such as mutant RAS (21). Passage through the bottleneck of p53-enforced growth arrest or apoptosis would seem to be a key step in almost any carcinogenic pathway.

Myc’s oncogenic impulse is mitigated by wildtype p53 but becomes unchecked when the tumor suppressor is absent or mutated (22). Moreover, wild-type p53 suppresses physiological *MYC* expression (23, 24). p53 transcriptionally upregulates its effector target genes while damping global transcription (24). In response to some genotoxic insults, p53 both prompts MYC degradation (25, 26) and decreases MYC expression (27). By decreasing MYC-driven global transcription amplification while simultaneously upregulating its own target effectors, p53 focuses cellular resources on the response to genotoxic stress.

Despite being closely regulated, there is no compensation mechanism that counteracts *MYC*-dosage insufficiency. *Myc*-haploinsufficient mice are smaller, but longer-lived and in many respects healthier than their wildtype litter mates (28). The cells and tissues of haplo-insufficient mice express half the wildtype levels of Myc, with no evidence of corrective upregulation (28, 29). The tumor incidence (per animal) of haplo-insufficient and wild-type animals is similar but, since the former are longer-lived, their tumor incidence per unit time is slightly reduced (28). Although pathologically increased MYC levels occur in many human cancers and enforced high-level Myc is tumorigenic in experimental models, it remains unclear whether low levels of Myc can sustain tumorigenesis or whether supra-physioloigcal levels of Myc are essential to amplify the programs of whichever oncogenes and mutant tumor suppressors are operating in cells on the path toward malignancy (22).

To explore the effect of reduced Myc levels on spontaneous oncogenesis, we compared tumorigenesis in *Trp53*^-/-^ mice (hereafter referred to as p53KO) that are either *Myc* wildtype (WT; *Myc*^+/+^) or haplo-insufficient (*Myc*^+/-^). *Myc*^+/-^;*p53KO* mice developed the same spectrum of tumors at similar frequencies to *Myc-*WT;*p53KO*, but tumor-free survival was extended by 50%.

Surprisingly, tumors from *Myc*^+/-^ versus WT mice expressed similar levels of Myc, indicating that during oncogenesis, the MYC-haploinsufficent gene dosage had either been compensated for, or bypassed. Whereas hemangiosarcomas arising in *Myc-WT;p53KO* mice were tetraploid, those arising in *Myc*^+/-^;*p53KO* were octoploid arguing that haploid Myc levels had been compensated to cross a threshold for increased Myc required in tumorigenesis. Thymic lymphomas from *Myc*^+/-^;*p53KO* mice also compensated MYC to the same high levels as *Myc-WT*;*p53KO* by another mechanism. These results indicate that Myc must be elevated for oncogenesis and that *Trp53*-loss alone is insufficient to drive Myc over this oncogenic threshold. Dropping Myc below the threshold may be a useful, achievable therapeutic goal in cancer patients.

## RESULTS

### *Myc* haploinsufficiency retards *p53KO* tumorigenesis

All of the tissues in *Myc*^+/-^ mice display half the levels of Myc protein as their WT counterparts (28, 29). Yet the incidence of cancer is not dissimilar between these mice, raising the question whether the difference in the physiological baseline of Myc was relevant to either the age of onset or progression rate of their tumors. If Myc is a transcriptional amplifier that intensifies the output of all already active cellular pathways (16, 18, 19), then oncogenic program(s) should be more forcefully driven in *Myc-WT* than in *Myc*^+/-^ cells. To test this idea, *p53*KO siblings that were either *Myc-WT* or *Myc*^+/-^ were monitored for tumor formation until either visible tumors were evident (hemangiosarcomas), or first signs of respiratory distress appeared (thymic lymphomas), at which point mice were euthanized and necropsied. On average, *Myc*^+/-^;*p53KO* mice lived ∼1.5x longer and up to twice as long as *Myc-WT*;*p53KO* offspring from the same cohort of breeders (**Figure 1**). The spectrum of tumors was similar irrespective of Myc-dose: most commonly hemangiosarcomas and thymic lymphomas; a small number of other tumor types including nodal and splenic lymphomas, germ cell tumors and adenocarcinomas, also occurred with lower frequency (Supplemental Table 1). This striking and highly significant difference in tumor-free survival between *Myc*^+/-^;*p53KO* and *Myc-WT*;*p53KO* mice for all major tumor types (**Figure 1A-D**), indicated that the pre-tumor basal level of Myc influences the pace of tumorigenesis. These experiments argue that haploid levels of Myc in pre-neoplastic tissues retard *p53KO*-driven tumorigenesis.

**Figure 1.**
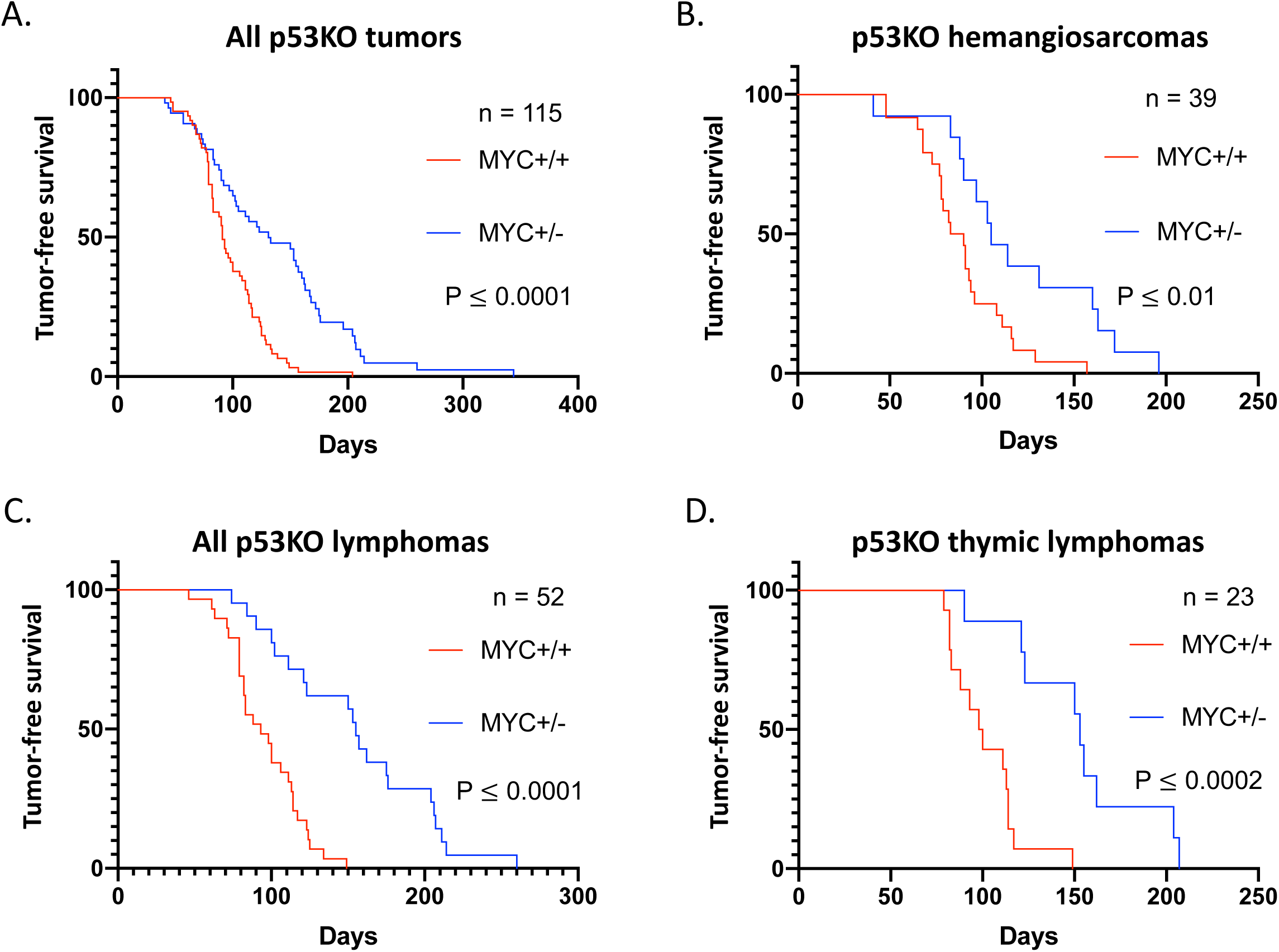
Extended tumor-free survival of p53KO mice that are haplo-insufficient for Myc gene dosage. Kaplan-Meier graphs of survival (days) until gross tumor appearance for all tumors, and for major tumor types occurring in p53KO, for Myc+/+ (red) and Myc+/-(blue). A. Survival curves for all tumor types (enumerated in Supplemental Table 1), including cases of incidental death without any evidence of tumor on necropsy (censored observations in Prism analysis). B. Survival curves for isolated hemangiosarcoma (no other concurrent tumors). C. Survival curves for all lymphomas (including thymic, nodal, splenic primaries, both with and without concurrent hemangioma). D. Survival curves for isolated thymic lymphoma (no other concurrent tumors). *p*-values for *Myc*-WT vs *Myc*^+/-^ Kaplan-Meier curves were calculated using Prism software.

### Hemangiosarcomas compensate for *Myc* haplo-insufficiency by gene dosage increase

To study the influence of Myc levels on tumorigenesis, we analyzed hemangiosarcomas, the most frequent tumor type occurring in the *p53KO* setting. If progressing more slowly, the *Myc*^+/-^;*p53KO* tumors might be expected to display half the Myc level of the *Myc-WT*;*p53KO* tumors (**Figure 1B**). Since solid tumors generally include both neoplastic cells and other components (stroma, vessels), Myc protein and RNA levels were evaluated *in situ* on tumor cell nuclei in histologic sections. Sections of *Myc*^+/-^;*p53KO* or *Myc-WT*;p*53KO* hemangiosarcomas were immuno-stained with anti-Myc and quantified (**Figure 2A**). Unexpectedly, the Myc protein levels of *Myc*^+/-^;*p53KO* and *Myc-WT*;*p53KO* tumor cells were similar. *Myc* mRNA levels quantified by RNA fluorescent *in situ* hybridization (RNA FISH, **Figure 2B**), likewise revealed no difference between *Myc*^+/-^;*p53KO* and *Myc-WT*;*p53KO* tumor cells. The equivalent Myc expression levels in *Myc*^+/-^ and *Myc-WT* tumors was unanticipated; in all published studies of cellular and organismal *Myc* haplo-insufficiency, such compensation to wild-type levels has not been reported in normal tissues. Moreover, if the tumors in the *Myc*^+/-^;*p53KO* versus *Myc-WT*;*p53KO* mice displayed no difference in Myc levels, then what might account for the longer survival of the *Myc*^+/-^;*p53KO* mice? We therefore further explored the mechanism of Myc dosage compensation in *Myc*^+/-^;*p53KO* hemangiosarcomas.

**Figure 2.**
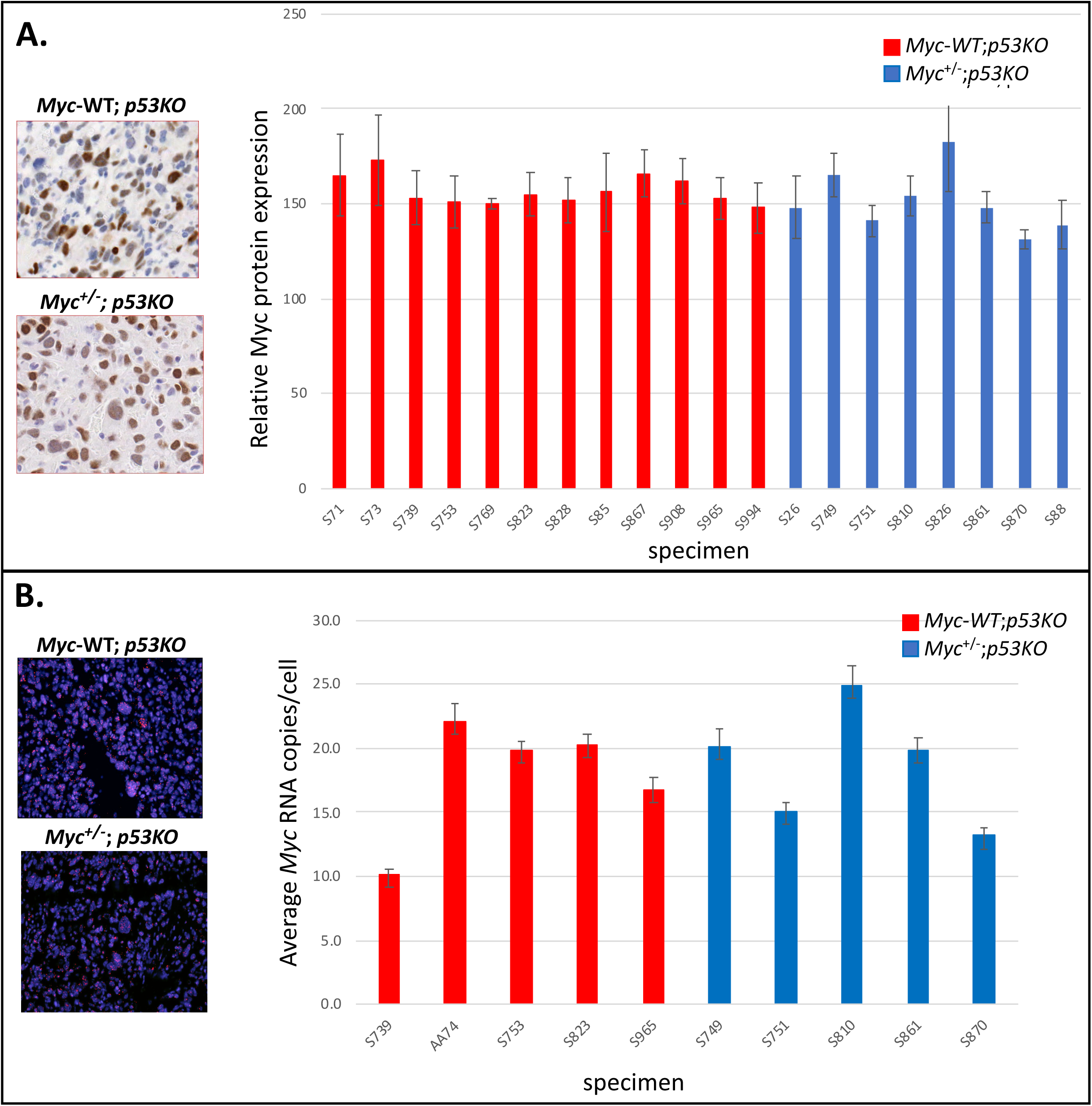
Comparable Myc RNA and protein expression levels in hemangiosarcomas from p53 KO mice that are either WT or haplo-insufficient for Myc dosage. A. Representative examples of Myc protein expression by immunohistochemical staining (IHC) and graphs of average Myc protein expression level in 12 independent cases of *Myc*^+/+^ and 8 independent cases of *Myc*^+/-^ p53KO-hemangiosarcomas. *Myc* protein was normalized relative to standard positive and negative controls included with each batch of automated immunohistochemistry. In all Figures, “specimen” identification numbers are indicated for independently occurring mouse tumors analyzed. (SEM indicated) B. Graphs of average *Myc* RNA copy number per nucleus from RNA FISH analysis in 4 independent cases of *Myc*^+/+^ and 6 independent cases of *Myc*^+/-^ p53KO-hemangiosarcomas. Bars indicate standard error of the mean (SEM, generated using Visiopharm software).

MYC over-expression in tumors frequently arises due to chromosomal rearrangements, including translocations and tandem gene amplification. Either of these events could compensate the difference in Myc levels present in the pre-malignant cells of *Myc-WT*;*p53KO* compared to *Myc^+/-^*;*p53KO* mice. *Myc* locus amplification is in fact a frequent feature of human angiosarcomas (30, 31). To evaluate *Myc* copy number and translocation-status in the *Myc^+/^*;*p53KO* and *Myc-WT*;*p53KO* hemangiosarcomas, DNA FISH was performed using dual color break-apart probes bracketing the *Myc* allele (**Figure 3A**). These red-green fusion signals co-localize at unrearranged *Myc* loci in histologic sections. Since the deleted coding segment of the knockout *Myc*-null allele resides entirely in-between these probes, the probes detect both, and do not discriminate between, the wild-type and knockout allele. No *Myc* locus rearrangements (separation of red and green probes) were identified in either *Myc-WT* or *Myc^+/-^* hemangiosarcoma cells; however, on average, the *Myc-WT* displayed ∼4 equally intense yellow spots per cell (overlap of the red and green breakaway probes), while the *Myc^+/-^* had ∼8 spots/cell (**Figure 3B, C**). Thus, the average number of *Myc*-alleles (both wild-type and knockout) in tumor cells from *Myc*-haplo-insufficient mice was twice that found in tumor cells from *Myc*^+/+^ mice.

**Figure 3.**
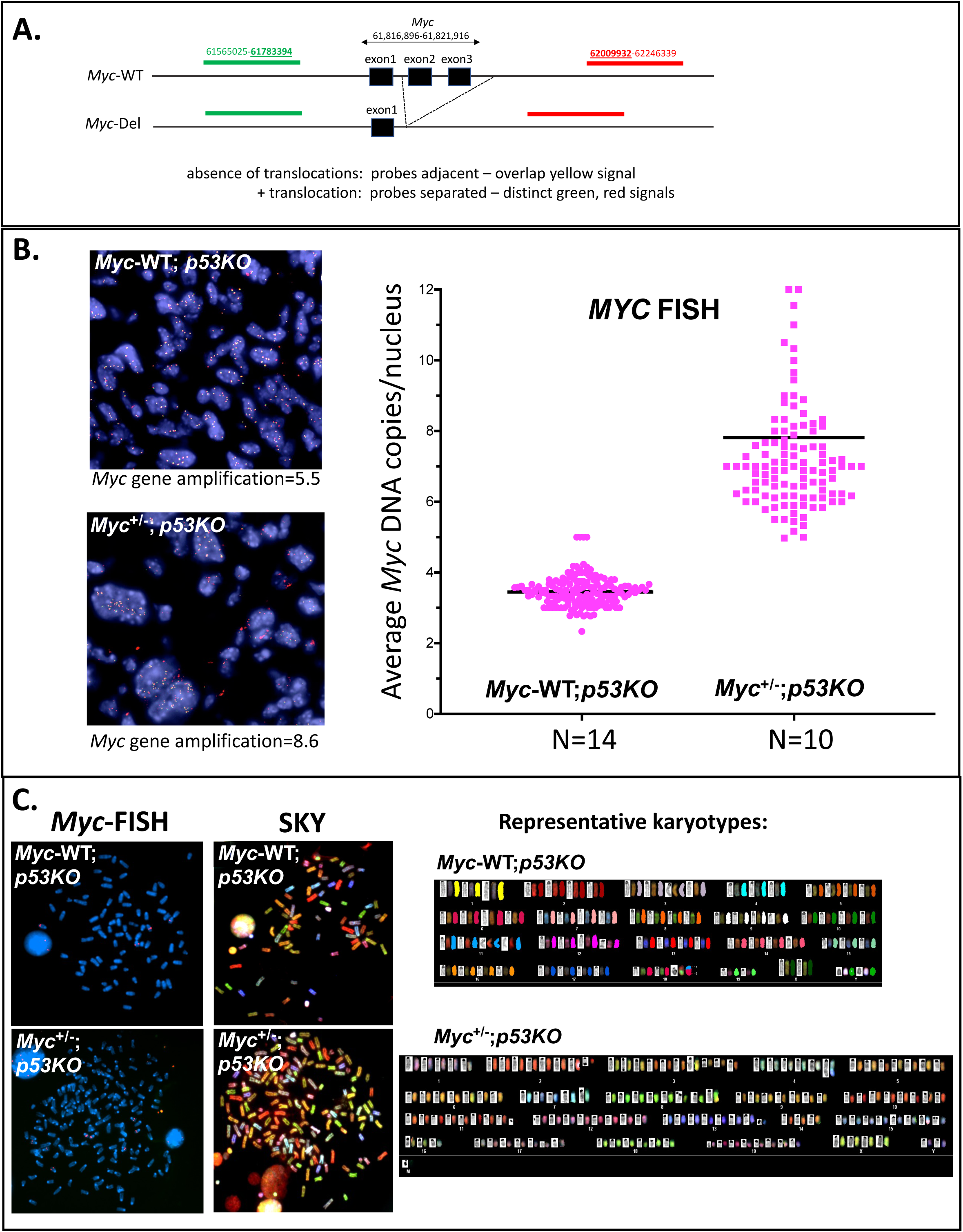
*Myc* gene dosage-compensation, augmentation and equalization by whole genome amplification in *Myc*^+/-^ compared with *Myc*^+/+^ hemangiosarcomas from p53KO mice. A. Schematic of *Myc* locus break apart BAC probe locations on mouse Chr15qD1 relative to WT and deleted *Myc* alleles (build mm9). B. Representative sections of *Myc* interphase DNA FISH in *Myc*^+/+^ and *Myc*^+/-^ p53KO-hemangiosarcomas and scatter plot of *Myc* DNA copy number in independent p53KO hemangiosarcomas from *Myc*^+/+^ (n=14 cases) and *Myc*^+/-^ (n=10 cases) mice. C. Representative examples of spectral karyotype analysis (SKY) in short term primary cell cultures derived from *Myc*^+/+^ (n=2 cases) and *Myc*^+/-^ (n=4 cases) p53KO hemangiosarcomas.

The karyotypes of the hemangiosarcomas arising in *Myc-WT*;*p53KO* versus *Myc*^+/-^;*p53KO* mice neatly accounts for the compensated *Myc*-gene dose of the latter (**Figure 3C**). Whereas, in the tetraploid (2/2) *Myc-WT* tumors all four alleles were functional, the *Myc*^+/-^ tumors were octoploid (4/4) and hence bore four native and four deleted (null) *Myc* alleles. Thus, all hemangiosarcomas possessed a net of 4 functional wild-type *Myc* alleles irrespective of genotype. Apparently, the Myc deficiency in the *Myc*^+/-^ mice selected for an extra round of whole genome duplication during hemangiosarcomagenesis. Furthermore, none of the tumors were diploid in *Myc* gene dose (n=24, **Figure 3B**), indicating that all hemangiosarcomas require supraphysiological levels of Myc achieved by endoreduplication.

### MYC-haploinsufficiency per se does not sponsor physiological hyperploidy

The octoploidy neatly compensating MYC-dose between *MYC*+/-versus *MYC*-*WT* hemangiosarcomas could result from either selective pressure on tumor cells to exceed an oncogenic threshold or from endoreduplication somehow provoked by reduced MYC levels, but unrelated to any neoplastic impulse. In fact, in the presence of mitotic inhibitors increasing *MYC*-levels promoted uncoupling of replication and cell division (endoreplication) resulting in tetraploid- and octaploidization. To ascertain the influence of MYC-haploinsufficiency on physiological polyploidization, the DNA content of *Myc^+/-^* and *Myc-WT* megakaryocytes was compared throughout thrombocytogenesis where physiologic cycles of replication with endomitosis result in hyperploidy (up to 128N), prior to platelet-yielding megakaryocyte fragmentation. Whereas either complete loss (32) or hyperactivity of Myc (33) each reduce megakaryocyte ploidy (reflecting, respectively, decreased DNA synthesis versus increased mitosis), *Myc*-haploinsufficiency only minimally affected megakaryocyte ploidy **(supplemental figure 1).** All of this evidence indicates that polyploidy seems unlikely to be an intrinsic and inevitable consequence of MYC haploinsufficiency. Most simply, stochastic or sporadically occuring octoploidy allows hemangiosarcoma precursors in *Myc^+/-^* mice to achieve the same Myc expression level needed to transcend the neoplastic threshold as in *Myc-WT*.

### Thymic lymphomas compensate for haplo-insufficient *Myc* by elevating RNA expression level

Is dosage compensation to the same supraphysiological Myc level in *Myc-WT*;*p53KO* and *Myc*^+/-^;*p53KO* tumors a feature unique to hemangiosarcomas, or do other tumors arising in this setting also demand supraphysiological Myc levels? Because thymic lymphomas were the second most frequently occurring tumor type in both *Myc-WT*;*p53KO* and *Myc*^+/-^;*p53KO* mice, we checked whether Myc expression in these tumors was also dosage compensated, and if Myc levels were also supraphysiologic in these tumors. Like all normal, non-neoplastic tissues in *Myc^+/^*^-^ mice, normal thymocytes express half the Myc RNA of *Myc-WT* animals and so are not dosage compensated. Comparison of non-cancerous juvenile thymus from *Myc*^+/-^;*p53KO* and *Myc-WT*;*p53KO* mice by *Myc*-RNA-FISH revealed that the *Myc*^+/-^ cells contained half the *Myc* RNA of the *Myc-WT* (**Figure 4**) as expected, and immunoblots revealed that haplo-insufficient Myc protein levels were also uncompensated to wild-type levels (**Figure 5B**). In contrast, comparing lymphomas arising in either the *Myc-WT;p53KO* or *Myc*^+/-^*;p53KO* mice, RNA FISH (Figure 5A), *Myc* RNA was equally elevated (**Figure 5A**). Concordantly, immunoblot and immunohistochemical staining revealed similar, markedly supraphysiological Myc protein levels in both *Myc*^+/-^;*p53KO* and *Myc-WT*;*p53KO* lymphomas (**Figure 5A,B**).

**Figure 4.**
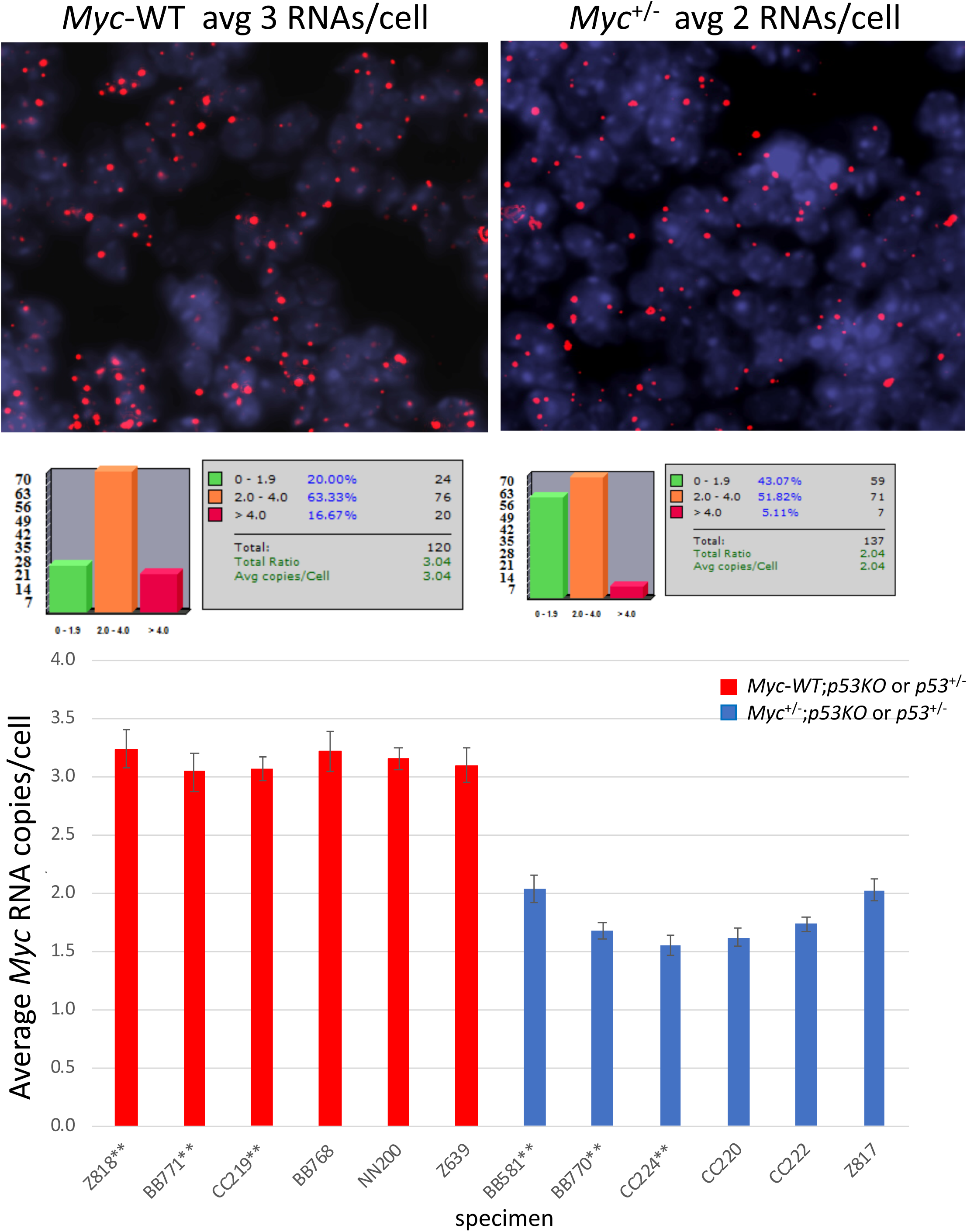
*Myc* RNA expression level in normal juvenile thymus from *Trp53*^+/-^ or *p53KO* mice are not *Myc* gene-dosage compensated. Representative sections of RNA FISH, relative *Myc* RNA copy calculations, and graphs of average RNA expression/lymphocytic nucleus in 6 normal juvenile thymuses from *Myc*^+/+^ (red) or *Myc*^+/-^ (blue) mice compared in *Trp53*^+/-^ and p53KO backgrounds. Bars indicate SEM, generated using Visiopharm software; (see also examples of normal thymic Myc protein in Fig. 5B). **indicate thymus from *Trp53*^+/-^ mice.

**Figure 5.**
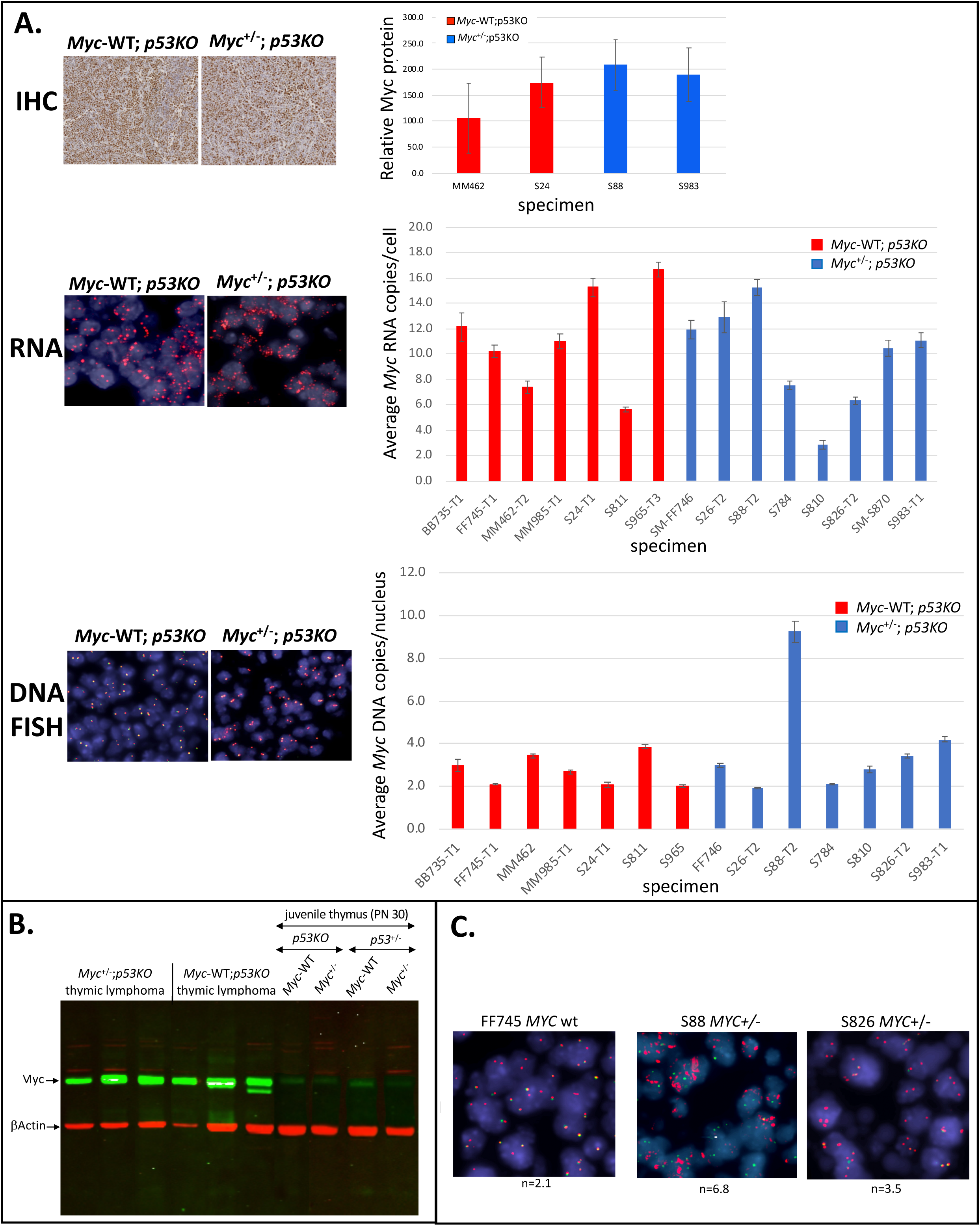
*Myc* RNA and protein expression are equalized without gene dosage-compensation between thymic lymphomas in *Myc*^+/-^ and *Myc*^+/+^ *p53KO* mice. A. Representative sections (left panels) and quantitation (right bar graphs) of *Myc* protein expression (as in Figure 2), RNA expression (RNA FISH), and DNA copy number (DNA FISH) per tumor cell. B. Immunoblot of representative *Myc*^+/-^ and *Myc*^+/+^ p53KO thymic lymphomas and normal (non-neoplastic) juvenile p53KO or *p53*^+/-^ thymuses, relative to β-actin loading control. C. *Myc* DNA FISH sections showing the single *Myc*^+/-^ thymic lymphoma in B. with *Myc* gene amplification (S88), compared to representative *Myc*^+/-^ and *Myc*^+/+^ lymphomas with minimal or no dosage changes. For all A panels, bars indicate standard error of the mean (SEM, generated using Visiopharm software).

However, in contrast to hemangiosarcoma, no single genomic process was discerned that could account for expression compensation in the *p53KO* lymphomas arising in the *Myc-*haplo-insufficient setting (**Figure 5A, B**). DNA FISH revealed no consistent *Myc* rearrangements, aneuploidy, polyploidy or *Myc*-gene copy number amplification in *p53KO* lymphomas from either *Myc*^+/-^ or *Myc-WT* mice (**Figure 5A, C**). In a single *Myc*^+/-^;*p53KO* lymphoma (of seven evaluated), significantly increased gene dose was detected, averaging ∼7 *Myc* alleles/cell distributed throughout tumor cell nuclei; these cells displayed a separation of the red and green “break-apart” *Myc* probe indicating chromosomal rearrangement, as well as amplification. Consequently, in the remaining *Myc*^+/-^ lymphomas, the net output of the single intact *Myc* allele matched the combined output of the two alleles in the *Myc-WT* lymphomas, in contrast to the situation in pre-neoplastic juvenile thymus. Apparently, multiple mechanisms may operate to upregulate *Myc* in thymic lymphomas, acting mainly at the transcriptional and post-transcriptional levels.

### Established thymic lymphomas demand elevated Myc levels

In experimental systems, tumors are “addicted” to Myc over-expression (34-37). In these systems, Myc is required not only transiently to push cells through a bottleneck early during oncogenesis but is necessary to maintain malignancy. We asked whether thymic lymphoma arising spontaneously in *p53KO* mice, without engineered *Myc* expression, would similarly be addicted to Myc expressed from its native locus. To address this question, we generated *p53KO* mice with a conditional, floxed *Myc* allele (*Myc^C/+^*) and a tamoxifen-activated CreER allele expressed ubiquitously from the Rosa locus (*Myc^C/+^;Rosa^CreER/+^*). Primary thymic lymphomas arising in either *Myc*^C/+;^*p53KO;Rosa*^CreER/+^ or *Myc-WT;p53KO;Rosa*^CreER/+^ mice were collected, dissociated into single cells and injected subcutaneously into the flanks of nude mice.

Allografted mice were treated either with tamoxifen to excise the floxed *Myc* allele (making the *Myc*^C/+^ tumors haploid in *Myc* dosage), or with vehicle alone, making the allografts derived from the same primary tumor either *Myc*-haploid or -diploid, respectively, depending on treatment (**Figure 6B,C**). Vehicle vs tamoxifen-treatment was also compared in *Myc*-WT tumors (**Figure 6A**) to assess the effects of tamoxifen alone on tumors harboring a CreER allele, but remaining *Myc*-diploid irrespective of tamoxifen treatment.

**Figure 6.**
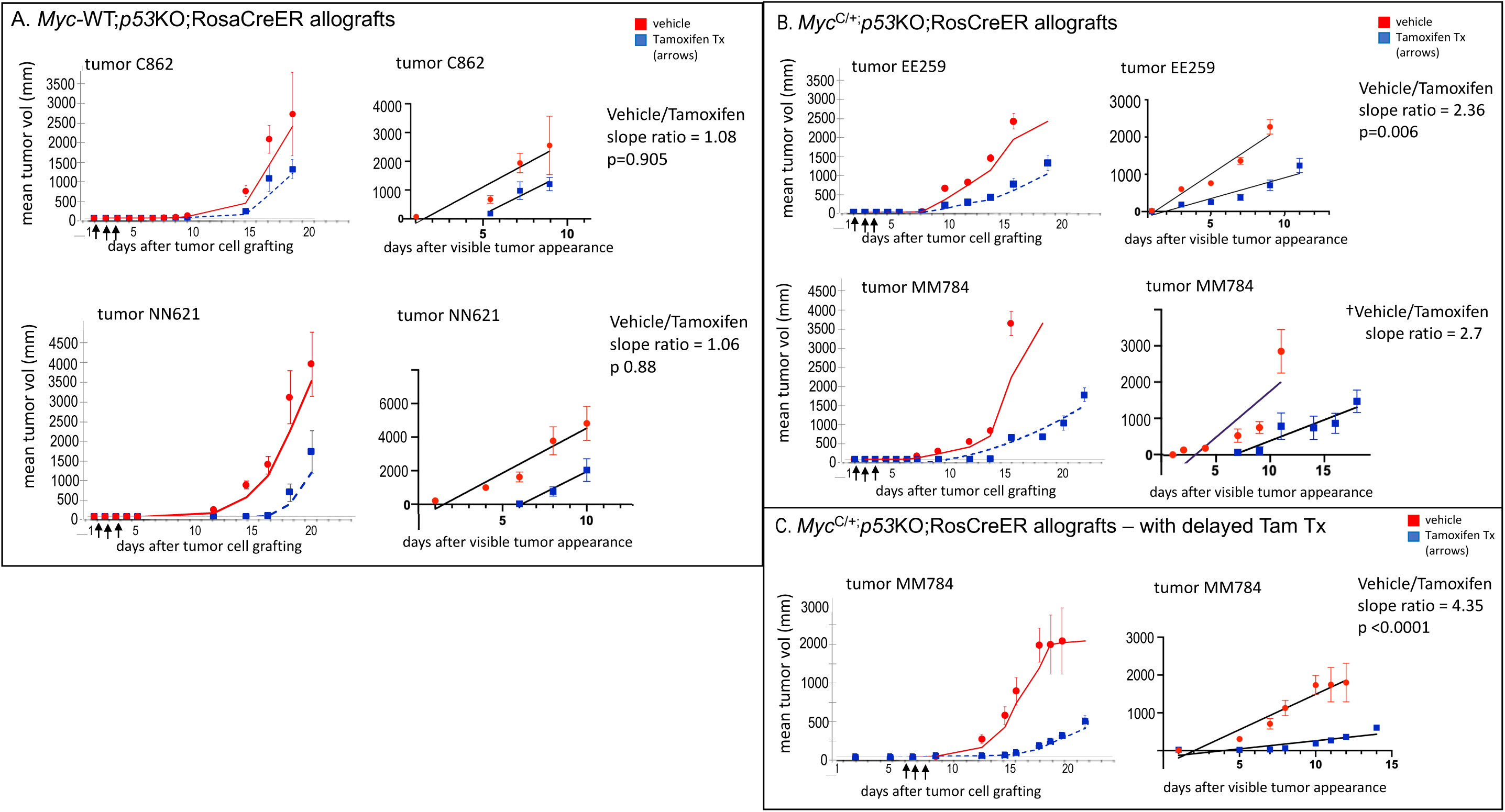
Acute reduction of *Myc* gene dosage in allografted, established *p53KO* thymic lymphomas impairs tumor growth. Dissociated cells from primary *Myc*^+/+^ (C862, NN621) or conditional *Myc*^C/+^ (EE259, MM784) *p53KO*;RosaCreER+ thymic lymphomas were injected subcutaneously in nude mice and treated in parallel with either tamoxifen (blue) or vehicle (red) for 3 consecutive days (arrows) beginning at either 1 day (A, B) or 7 days (C) after allografting. Mean tumor sizes were determined at each time point (raw data plots shown in **Supplemental Figure 2**) and best fit curve calculated using moving averages for vehicle and tamoxifen-treated groups (left panel graphs, Excel). Mean tumor sizes during the linear growth phase (after tumor became visibly apparent) were re-plotted to compare slopes (growth rates) and statistical significance (p) of slope differences between vehicle and tamoxifen-treated groups (right panel graphs, calculated with Prism 8 software). Error bars indicate standard error of the mean. A. Mean tumor allograft growth (mm^3^) for *Myc*-WT C862 and NN621 primary tumors, treated with vehicle or tamoxifen beginning 1 day after allograft. Growth rates (slopes) during the linear phase for vehicle and tamoxifen-treated are not statistically different. B. Mean tumor allograft growth (mm^3^) for *Myc*^C/+^ EE259 and MM784 primary tumors, treated with vehicle or tamoxifen beginning 1 day after allograft. Growth rates (slopes) during the linear phase for vehicle and tamoxifen-treated are significantly different (slope ratio 2.36, p 0.006) for EE259, indicating that acute *Myc* dosage reduction (tamoxifen treated) impaired tumor growth relative to control. Growth rates (slopes) during the linear phase for vehicle and tamoxifen-treated for MM784† also suggested that acute *Myc* dosage reduction (tamoxifen treated) impaired tumor growth relative to control (slope ratio est. >2.7), but p value could not be reliably determined because vehicle-treated tumors grew so rapidly that insufficient time points were available for linear regression analysis. See also C below. C. Mean tumor allograft growth (mm^3^) for *Myc*^C/+^ MM784 primary tumor, treated with vehicle or tamoxifen beginning after a delay of 7 days after allograft. Growth rates (slopes) during the linear phase for vehicle and tamoxifen-treated are significantly different (slope ratio 4.35, p <0.0001), indicating that acute *Myc* dosage reduction (tamoxifen treated) impaired tumor growth relative to control, even when recombination was delayed for a week to allow greater tumor engraftment.

Cre-induced apoptosis has been observed in the context of induced Cre-driven recombination (38, 39), although loss of *Trp53* (*p53KO*) mitigates Cre activity-induced apoptosis *in vivo* in mice (39). Nevertheless, apoptosis following tamoxifen treatment in *p53KO* thymic lymphomas harboring an engineered CreER allele has been previously reported (38). Consistent with this finding, both *Myc*-WT (**Figure 6A**) and *Myc*^C/+^ (**Figure 6B,C**) -*p53KO* lymphoma allografts displayed a variable delay in emergence of detectable tumors in the tamoxifen-treated compared to control groups for the same primary tumor allografts. However, if the tumor population size is simply reduced by a pulse of Cre-induced apoptosis, then upon tamoxifen withdrawal, once tumors appear, similar growth rates would be expected in both groups if tamoxifen-treatment acts only to reduce the initial tumor cell number. Indeed, once linear-growth phase resumed, the growth rates (slopes) for tamoxifen- and vehicle-treated groups in *Myc*-WT tumors were very similar (**Figure 6A**). But in the case of *Myc*^C/+^ lymphomas, a persistent, highly reduced growth rate was apparent for the same tumor after *Myc* gene dosage reduction (lower slope, **Figure 6B**); this differential effect of the induced *Myc* haplo-insufficiency remained evident even when tamoxifen treatment was delayed for a week after allografting to ensure efficient tumor engraftment (**Figure 6C**). Thus, the output of both *Myc* alleles must be maintained in *Myc*^C/+^;*p53KO* lymphoma to manifest full malignant potential. Even an incremental reduction in Myc tempered the aggressiveness of already established tumors.

### Do MYC levels correlate with DNA damage in hemangiosarcomas or thymic lymphomas in *MYC+/-* and *MYC*-WT, *p53KO* mice?

In the absence of *Trp53*, Myc levels are supraphysiologic in hemangiosarcomas and thymic lymphomas arising in both *Myc*^+/-^ and *Myc-*WT mice. The well-known antagonism between high levels of MYC and wild-type TP53 made us wonder whether, in the context of guarding the genome, TP53 directly or indirectly surveils MYC. Pathologically high MYC over-drives transcription and replication, provoking DNA-damage. This damage would be expected to elicit the TP53-meditated growth-arrest or killing of rogue, hyper-MYC cells. To assess DNA damage in *Myc*^+/-^ and *Myc-WT* tumors in mice, histological sections were probed for γH2AX, the standard marker for DNA damage (40), and for MYC and analyzed using quantitative immunohistochemistry (**Supplemental Figure 2A-C**). Across different hemangiosarcomas in *p53KO* mice, MYC and γH2AX levels correlated well, whether from *Myc*^+/-^ or from *Myc-WT* tumors, and the ranges of both MYC and γH2AX were fairly narrow (**Supplemental Figure 2A, 3A**). In contrast, in thymic lymphomas, γH2AX and MYC levels showed weaker correlation and much greater variability for both genotypes (**Supplemental Figure 2B, 3A**).

This case-to-case variation and weaker correlation of MYC and γH2AX levels in lymphomas, as compared to hemangiosarcomas, paralleled greater variation in overall RNA, protein, and DNA status (genome rearrangements, etc.) **(see Figure 5).** Notably, in lymphomas, γH2AX and MYC scores were 10X more variable than in hemangiosarcomas. Thus, thymic lymphomas are more heterogeneous than hemangiosarcomas, with respect to both MYC levels and DNA-damage, in the absence of p53 function.

The good correlation between MYC and γH2AX in hemangiosarcomas suggested that some pervasive MYC-related process might be stressing the genome throughout the tumor cell population, not just in sporadic cells. The recent discovery that Myc combines and catalytically activates topoisomerases 1 and 2 into a “topoisome” complex raised the possibility that these Myc-activated topoisomerases might acutely increase baseline DNA damage (41). Topoisomerase activity is intrinsically mutagenic (42, 43) and topoisomerase-poisons damage DNA and trigger a vigorous p53-response (41).

To test whether excess Myc is acutely genotoxic, inducible HOMycER12 cells (HO15.19 rat1a *Myc^-/-^;*Tg-MycER (44)) were treated either with tamoxifen or vehicle for fifteen minutes, and then treated for fifteen minutes with or without topoisomerase poisons prior to staining for γH2AX (**Figure 7A**). Surprisingly, the γH2AX signal provoked by activating MycER with tamoxifen alone was comparable to that elicited by the combined treatment with topoisomerase poisons camptothecin (CPT) and etoposide (ETO) (**Figure 7B, C**). Together, combined tamoxifen and CPT/ETO acted synergistically to increase the γH2AX signal to levels 3-fold greater than their sum. Vehicle treatment alone (**Figure 7B, C**) elicited no γH2AX signal. The rapidity of this γH2AX response made it unlikely to represent an indirect consequence of either transcription- or replication-generated stress (45, 46) following Myc-upregulation. The rapid onset and profound synergy between tamoxifen and CPT/ETO treatment strongly indicates a linkage between Myc activation and topoisomerase-mediated DNA damage. We recently found that at high levels, Myc assembles with topoisomerases 1 and 2 into a “topoisome” complex and stimulates their activities (47). By responding directly and immediately to topoisome-provoked DNA damage, p53 would cull tissues of cells that harbor incipient pathologic levels of Myc (**Figure 7D**).

**Figure 7.**
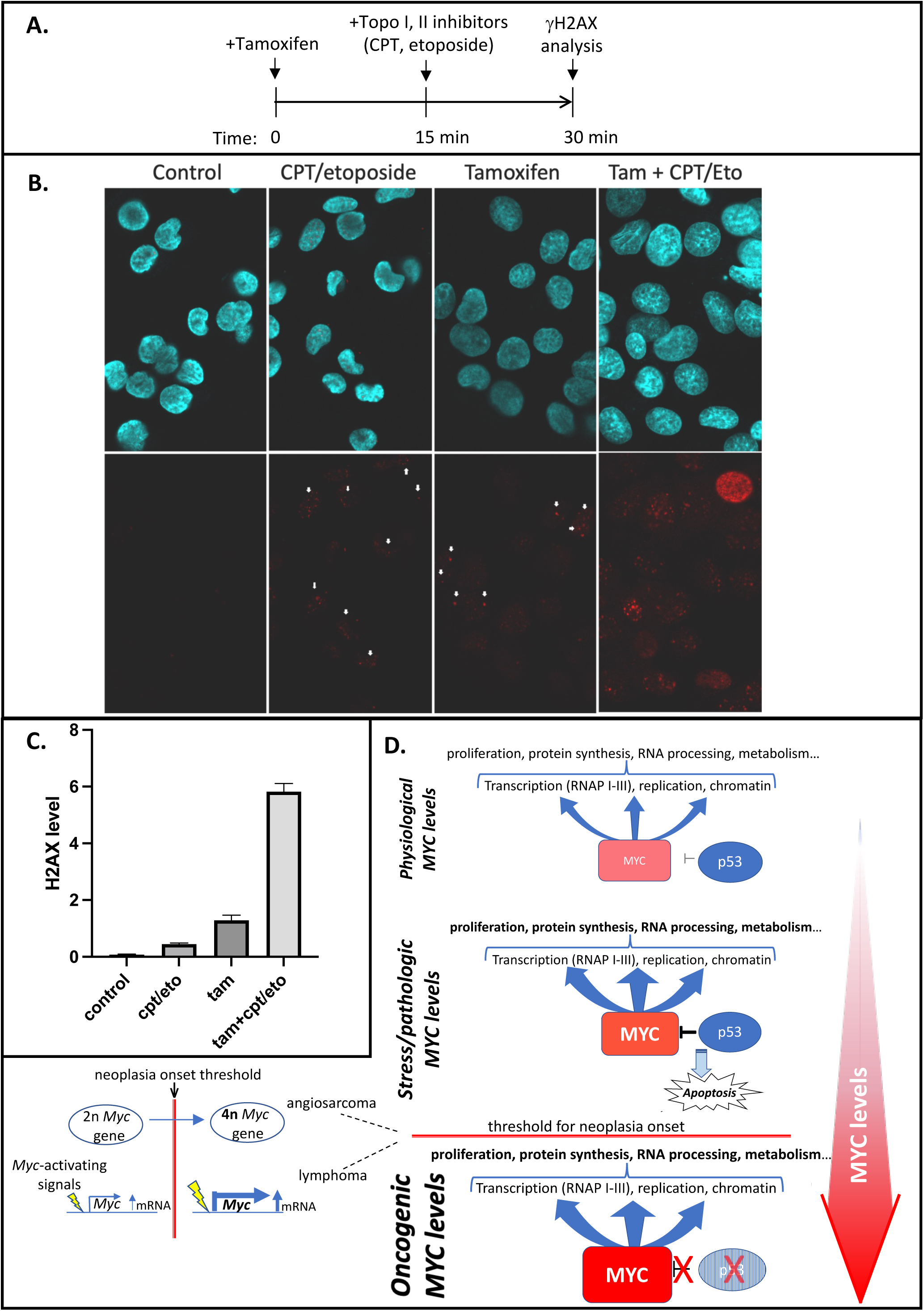
p53 rapidly senses high-Myc level in cells via DNA damage induced by Myc-topoisomerase complexes. A. Schematic timeline for treatment of cells, HOMycER12(44), expressing tamoxifen-inducible MycER with topoisomerase 1 (CPT) and 2 (etoposide) inhibitors, and 4OH-tamoxifen. B. MycER-expressing cells (HOMycER12) were treated as outlined in A, and immunofluorescence stained with anti-γH2AX to assay acute DNA damage at 30 minutes of MycER activation. Representative fields are shown. Arrows indicate examples of increased nuclear γH2AX fluorescence in CPT/etoposide and Tamoxifen treated cells. Increased fluorescence with Tam + CPT/Eto is unmarked. C. Graph of of experiment in B; error bars indicate SEM, n=4 independent biological replicates. P-values between all pairs of treatments ≤ 0.003 (standard 2-sided T test). D. Model for physiologic and oncogenic Myc roles. Top--In normal cells Myc levels are low and the loss of p53 provides an insufficient oncogenic impulse. Middle--As Myc levels rise p53 checks oncogenesis by 1) downregulating *Myc* transcription oncogenesis and 2) provoking apoptosis when Myc levels reach pathologic levels. Bottom— without p53, Myc levels may sustain pathologic levels above and oncogenic threshold. Left--Myc levels may transcend this threshold through a variety of molecular mechanisms such as the polyploidy that elevates *Myc* copy number in angiosarcomas, hyper-signaling to *Myc* causing transcriptional overdrive in lymphomas or by other mechanisms that increase *Myc* gene dose or mRNA levels or that stabilize Myc protein. See text for details.

### Spontaneous tumors in *Myc* haplo-insufficient mice also compensate for reduced *Myc* gene dosage

The similar Myc expression levels in the *Myc*^+/-^;*p53KO* and *Myc-WT*;*p53KO* hemangiosarcomas and in thymic lymphomas suggested that equivalent but supraphysiological levels of Myc are needed for tumorigenesis in both genotypes. Was this requirement unique to the *p53KO* mice or is *Myc* dosage compensation a general feature of tumorigenesis in *Myc^+/-^* versus *Myc-WT* mice irrespective of germline *Trp53* status? Sedivy and colleagues have previously characterized C57BL/6 mice carrying the same germline *Myc*^+/-^ null allele used here and have compared the long-term natural history of *Myc-WT* versus *Myc*^+/-^ mice with respect to the biology and pathology of aging (28). The major cause of death among both genotypes was cancer (∼80%) and among these tumors ∼80% were B-cell lymphomas. We compared *Myc* expression using RNA-FISH in sections of five lymphomas arising in *Myc-WT* versus five lymphomas occurring in *Myc*^+/-^ mice (**Supplemental Figure 5**). Comparable *Myc* mRNA levels were present in both groups indicating that tumorigenesis effaced the pre-neoplastic two-fold difference in *Myc* expression seen in the normal tissues of *Myc*^+/-^ versus *Myc-WT* mice.

## DISCUSSION

Together, *TP53* loss or malfunction and MYC over-expression occur in most human cancers. Here, we asked if altered endogenous Myc levels impact the tumorigenicity of tumor suppressor loss. Indeed, reducing germline *Myc* by 50% prolonged survival across the full spectrum of tumor types in mice lacking *Trp53,* indicating that Myc acts downstream of, or in parallel with, any cell-type specific oncogenic pathways. Although halving the basal Myc-dose in *p53KO* mice substantially extended tumor-free lifespan, animals inevitably succumbed to cancer (>90%) suggesting that extra Myc accelerates a pathologic trajectory programmed by the germline loss of *Trp53*. By increasing all transcription, Myc might provide a generalized oncogenic impulse that augments any active tumorigenic process independent of tissue type. In this scenario the pathogenetic processes are the same in *Myc*^+/-^ and *Myc-WT* cells, but are slower-paced in the former. Not accounted for in this scenario is the rise of Myc in haplo-insufficient tumors to the same elevated levels as their wild-type counterparts. As a group, thymic lymphomas made more Myc than hemangiosarcomas (by immunohistochemistry), yet within each tumor type, haplo-insufficient and wild-type Myc expression reached parity. Hence, both baseline and oncogenic Myc expression appear to be tissue-specific.

In hemangiosarcomas, the increase in Myc is paralleled by an increase in γH2AX (scored by immunohistochemistry) suggesting that Myc provokes DNA damage either directly via increased topoisome activity, or indirectly by creating transcription- and replication-stress, albeit within a very short time-frame (15 min). In *Trp53* wild-type cells, such damage would trigger senescence or death thereby culling high-Myc cells to guard against oncogenic transformation (**Figure 7D**, top panel); in *p53KO* cells the high-Myc cells survive to sponsor genotoxic-stress that impels further tumorigenesis (**Figure 7D**, middle, bottom panels) (7, 23). In the case of *p53KO* hemangiosarcomas, the mechanism that upregulates Myc is clear; whole genome duplication (2nè4n) (**Figure 7D**, bottom left). In the Myc-WT tumors, one round of endo-reduplication yielded tetraploid cells with four intact copies of *Myc*. In the *Myc*^+/-^ tumors, the first round of endoreduplication to tetraploidy restored *Myc* to its normal diploid level; an amount of *Myc* apparently insufficient to support hemangiosarcomagenesis. A second round of endoreduplication to octoploidy in the *Myc*^+/-^ background results in four functional *Myc* alleles—the same number as in *Myc*-*WT* tumors. These results indicate that Myc is required at twice its WT level during hemangiosarcomagenesis. The tetraploidization of *Myc-WT* tumors versus octoploidization of *Myc*^+/-^ tumors (**Figure 3B, C**), argues that a requirement for *Myc*-gain is the principal driver of endo-reduplication (although Myc-haploinsufficiency per se does not drive physiologic endoreplication). Similarly, in a mouse model of acute promyelocytic leukemia (APL), trisomy of the *Myc*-bearing chromosome 15 is suppressed when extra Myc is supplied via retroviral expression (48). In the case of murine APL model, a *Myc*-gain of 50% seems to be a sufficient oncogenic driver of trisomy. Notably trisomy of the *MYC*-bearing human chromosome 8 is also frequent in APL (48). Thus, modest changes in Myc levels can profoundly impact tumorigenicity. Though tetraploidization in tumors arising in *Trp53*-null conditions has been noted previously, the biological driver of this event was not definitively identified (49-51). The current results suggest that a requirement for a gain of Myc may be such a driver.

Between *Myc*^+/-^;*p53KO* and *Myc-WT*;*p53KO* thymic lymphomas, Myc levels also were also elevated and equalized. However, unlike hemangiosarcomas, the magnitude of the increased expression, relative to non-transformed wild-type lymphocytes, was far greater and more variable than two-fold. Because chromosomal rearrangements and *Myc* gene amplification were infrequent, we surmise that elevated Myc is due to pathologic trans-activating signals delivered to the *Myc* locus (**Figure 7D**, bottom left). Notch pathway activation of *Myc* occurs frequently in thymic T-cell lymphomas. Defective Ikaros-repression in these lymphomas may upregulate Notch targets (52). However, such trans-activation would be expected to drive both alleles in *Myc-WT* cells, but only one in *Myc*^+/-^ cells, leaving equalization of Myc expression unexplained. Yet such two-fold effects may be overwhelmed by variable high-level activation of Myc expression. Additionally, in lymphomas, total Myc output may be limited by Myc-autorepression as first observed in Burkitt lymphoma, where enforced expression of the translocated *MYC* allele results in silencing or near silencing of the normal allele (53, 54). If negative auto-feedback in *trans* limits Myc expression by direct or indirect mechanisms, though *Myc*^+/-^ cells would initially lag wild-type Myc levels, as expression rises to the levels seen in lymphomas, both genotypes would converge to the same level. Autorepression would trend to the same limit irrespective of gene copy number. Stabilization of MYC protein in thymic lymphomas by mutations in E3-ligases (that otherwise target MYC degradation) also likely contributes to a more variable ratio of *MYC* RNA to protein level than is seen in hemangiosarcomas.

Weaker correlation of γH2AX and MYC in thymic lymphomas than in hemangiosarcomas likely reflects the special biology of lymphocytes. Superimposed on the DNA-damage directly or indirectly attributable to MYC, is damage due to ongoing expression of genome-modifying enzymes that expand T-cell receptor repertoire during lymphocyte development (i.e. Rags 1&2, AID) (55).

Our results also support the notion that a threshold for Myc that must be exceeded to sponsor tumor growth (22, 34, 56, 57). Deleting one conditional *Myc* allele in thymic lymphomas allografted into nude mice initially retarded tumor growth rate, indicating a sustained requirement for high-level Myc. This observation is similar to the demonstration that tumors arising in the setting of experimentally enforced Myc expression are oncogene “addicted”, with the difference being that elevated Myc expression levels were not enforced in the current study. Acutely decreasing Myc levels below a “threshold” in either situation interrupts neoplasia until the surviving cells can re-establish a mechanism to achieve Myc upregulation (34). This finding appears to be so consistent, at least in the spectrum of *p53KO*-driven tumors, that it suggests that restraining Myc levels, rather than guarding against the effects of unrepaired random DNA damage per se, constitutes the major p53 tumor suppressor function. The immediate and delayed DNA-damage consequent to elevated Myc enables p53 to continuously purge any cells that exceed Myc’s oncogenic threshold.

That different tumors exploit different strategies to upregulate Myc is likely a reflection of the complexity of how *Myc* expression is regulated. Transcribed from multiple promoters driven by a broad panel of enhancers and super-enhancers strewn across several megabases and regulated at every possible stage of macromolecular synthesis and degradation (11, 13, 58), *Myc* is held to different setpoints and tolerances in different tissues and cell types. Myc function may also be modulated by negatively acting members of the Myc network that titrate out interaction with its partner Max (59, 60). During oncogenesis, incipient tumors must defeat whichever safeguards restrict *MYC* expression in whatever setting they arise. Thus, universal therapeutic schemes to thwart pathological regulation of *MYC* expression may prove elusive and such strategies may need to be individually tailored to intercept tumor-specific *MYC*-upregulating pathways.

Conversely, intercepting/modulating MYC function may have general utility in a broad range of tumors regardless of pathogenic drivers. All the tumors seen in these experiments expressed supraphysiological Myc levels, albeit sometimes only modestly, including the spontaneous tumors arising in *Trp53-WT* mice. If this is general, it would argue that there is no efficient substitute or work-around for loss of endogenous Myc function in neoplasia. Myc elevation above a physiologic threshold maybe a prerequisite for tumorigenesis irrespective of *Trp53* status. Through the years, attempts to complement Myc-deficiency for tumorigenesis by supplying other oncogenes and candidate Myc-targets singly or in combination have all failed (other than with *Myc*-family members) (61). The sensitivity of tumorigenesis and tumor progression to the relatively small differences between haplo-insufficient and wild-type Myc levels, or between diploid and tetraploid cells to sponsor hemangiosarcomagenesis, argues that partial inhibition of Myc function may suffice to block or significantly retard disease progression in most cancers. In the setting of genetic *TP53*-loss (Li-Fraumeni syndrome)(62), even a partial antagonism/inhibition of MYC activity would be expected to moderate tumor incidence/onset and temper tumor aggressiveness. Besides such prophylaxis, partial inhibition of MYC activity in an adjuvant setting might help to attenuate tumor recurrence or could enhance the efficacy of combination chemotherapy.

## METHODS

### Mouse lines, tumors collection, allografts and tamoxifen treatment

All animal studies were carried out according to the ethical guidelines of the Institutional Animal Care and Use Committee (IACUC) at NCI-Frederick under protocol #ASP-17-405. Conditional, floxed *Trp53*^C/C^ (63) and *cMyc*^C/C^ (64) alleles, and tamoxifen-dependent *Rosa*CreER (65) mouse lines have all been described previously. *c-Myc*-deleted (*Myc*^+/-^) and *Trp53*-deleted (*Trp53*^+/-^) mice were each generated by crossing *Myc*^C/C^ or *Trp53*^C/C^ males with *Prrx1*Cre females to produce germ-line recombination (66). All *Myc*^+/-^ and *Myc*^+/+^ mice for tumor studies were subsequently generated from the same sets of breeders in *Myc*^+/-^;*Trp53*^+/-^ x *Trp53*^+/-^ crosses to minimize genetic background variation. *Trp53*^-/-^ (p53KO) offspring were monitored regularly (at least 3-4 times/week) for evidence of visible tumor development (usually hemangiosarcomas) and for any symptoms of illness (particularly dyspnea/distress for the other common tumor type, thymic lymphoma, which presents with shortness of breath). Mice with visible tumors, dyspnea or other symptoms were euthanized immediately, necropsied and tissues evaluated. Tumor histology was reviewed by at least 2 pathologists for diagnosis.

For allograft experiments to examine the consequences of *Myc* gene dosage reduction in established thymic lymphomas, primary *Myc*^C/+^;*Trp53*^-/-^;RosaCreER tumors were dissociated and single cells recovered by passing tissue through a 70um nylon mesh cell strainer with sterile Dulbecco’s PBS and cryopreserved (20% FCS, 10%DMSO). To generate a large number of tumor cells with high viability, cryopreserved cells were injected subcutaneously into the flank of nude mice (1-2x10^6^ cells). When an aggregate tumor size of 2 cc was reached, the primary allografts were collected (yielding 1-2 x 10^8^ cells) and freshly dissociated lymphoma cells were immediately re-injected subcutaneously into the flank of new recipient nude mice (ranging from 0.5 - 5 x 10^6^ cells in 0.1cc sterile PBS, indicated in text). Allografted nude mice then received intraperitoneal injections of 2mg tamoxifen in sterile vegetable oil (67) (0.2cc, Sigma, T-5648) or vehicle alone for 2-3 consecutive days after tumor injection, as indicated in text. Tumor injection sites were monitored daily for growth and tumor measurements were taken with calipers (length, width, height), approximately every 48 hrs. Animals were euthanized and tumor collected when an aggregate size of 2cc was reached, or earlier if any tumor erosion or signs of distress occurred.

### Detection of *Myc* gene copy number alterations and structural rearrangements by FISH on metaphase chromosomes and FFPE mouse tumors

The *Myc* gene copy number changes and rearrangements in the tumors were analyzed using Fluorescence *In Situ* Hybridization (FISH). The *Myc* Break-Apart FISH probe was designed using BAC DNA probe contigs RP23-7F11, RP23-376L10, RP23-130M7 labeled with Spectrum Green, covering myc 5’ upstream region (586kb) and RP23-137B1, RP23-393C9, RP23-450J14 labeled with Spectrum Orange Fluorescence covering *Myc*-3’ downstream region (584kb) separated by a 699kb gap between 5’ and 3’ probes. BAC clones were purchased from Empire Genomics (Buffalo, NY). FISH assays were performed on metaphase chromosomes or 5-micron Formalin-Fixed-Paraffin-Embedded (FFPE) tumor sections using a standardized protocol with slight modifications (68). In brief, 5μm thick sections of FFPE tissue blocks were de-paraffinized, rehydrated, and antigen retrieval performed with IHC-Tek Epitope Retrieval Solution (IHC World, Ellicott City, MD) with steaming for 25 min. After cooling, slides were treated with 50 μg/ml pepsin at 37°C, rinsed in PBS solution followed by dehydration in ethanol series. Co-denaturation of the probe and target DNA at 73°C in HYBrite (Abbott Molecular, Chicago, IL) for 5 min was followed by overnight hybridization at 37°C. Slides were then washed at 72°C in 0.4X SSC/0.3% Tween-20 for 2 min and cooled in 2X SSC/0.1% Tween-20 at room temperature for 1 min. The slides were counterstained, mounted with DAPI/Antifade (Vector Laboratories, Burlingame, CA) and analyzed on the BioView Duet-3 fluorescent scanning station (BioView, Billerica, MA) using 63X-oil objective and DAPI/FITC/Rhodamine single band pass filters (Semrock, Rochester, NY. A minimum of 100 tumor cell nuclei were scored for each specimen.

### Metaphase chromosome preparation from hemangiosarcoma established cell lines for Spectral Karyotyping

Freshly-collected hemagiosarcomas were minced and multiple small cell clusters were plated on T75 flasks in Dulbecco’s modified Eagle’s medium (DMEM) with 20% FBS, glutamine, and antibiotics. After outgrowth, cell colonies were trypsinized with 0.25% trypsin in bulk and passaged upon confluence. Metaphase chromosome harvesting was done using standard procedures (68). Cells were arrested with Colcemid 0.05 μg/ml for 3 hours in log phase, trypsinized, and subjected to hypotonic treatment with 75 mM KCl for 30 min at room temperature, followed by fixation in cold fresh methanol-acetic acid (3:1) for 20min followed by three changes with fresh fixative. Metaphase spreads were prepared by dropping 15ul of cell suspension on clean positively-charged slides. Slides were air-dried and analyzed by FISH as above. Spectral karyotyping (SKY) was performed using a mouse SKY kit from Applied Spectral Imaging (ASI, Carlsbad, CA) on metaphase chromosomes from short-term cultured mouse hemangiosarcoma tumor cells according to the manufacturer’s protocol.

### RNA *In Situ* hybridization

RNAscope analysis for *Myc* RNA expression was done using mouse *Myc* RNAscope probe and kit from Advanced Cell Diagnostics (ACD). Mouse ACD target probe was Mm-Myc (Cat#. 413451-C2). Slide pre-treatment and hybridization steps were performed according to the manufacturer’s protocol (https://acdbio.com/technical-support/user-manuals).

Fluorescence Imaging was done on the BioView Duet-3 fluorescent scanning station (BioView, Billerica, MA) using 63X-oil objective and DAPI/FITC/Rhodamine single band pass filters (Semrock, Rochester, NY. At least, 100 tumor cell nuclei were scored for each specimen. For FISH analysis *Myc* Break-Apart program was used for signal pattern analysis and quantification. The cut-off level for a tumor to be considered positive for *Myc* rearrangement (resulting in split red/green signals) was 10%. Quantitative RNA analysis was done using the HER2/neu amplification BioView software.

### Immunohistochemistry of Myc protein in hemangiosarcomas and thymic lymphomas

Myc protein expression analysis was performed using rabbit MAB EP121 (Epitomics, Cat#AC-0116), 1:200 dilution with high pH HIER. Image scanning and quantitative analysis was done using NanoZoomer scanner and Visiopharm (Visiopharm, Westminster, CO) image analysis software. Nuclear *Myc* protein staining was normalized relative to standard positive and negative controls included with each batch of automated immunohistochemistry.

### Immunoblot analysis of Myc protein in normal juvenile thymus and thymic lymphomas

For western immunoblot analysis, normal juvenile thymuses of 28-30 day old *Trp53*^-/-^ or *Myc*^+/-^;*Trp53*^-/-^ mice or primary thymic tumors were isolated, minced and homogenized in 1% SDS, 50mM Tris PH7.5, with protease inhibitors and AEBSF added. For all normal and tumor tissues analyzed, histologic sections were also examined to confirm either absence of cryptic early lymphoma (postnatal normal thymus) or to confirm lymphoma. Lysate proteins were electrophoresed in NuPAGE 10% Bis-Tris protein gels (Invitrogen), transferred to nitrocellulose membranes and probed with either affinity-purified monoclonal rabbit anti-Myc (Abcam ab32072; 1:10,000) and mouse anti-beta-actin (Abcam ab184576; 1:10,000), and visualized with fluorescent secondary antibodies (LI-COR IRDye 800CW, 926-32211, anti-rabbit green; and with #680RD, #926-68072, anti-mouse red) using LI-COR Odyssey CLx.

### Analysis of ploidy in megakaryocyte/thrombocytic lineage of bone marrow progenitors from Myc-WT and Myc+/-mice

To analyze the distribution of ploidy in megakaryocyte (MK) subpopulations at different maturation stages (I – V), bone marrow cells were flushed from femur and tibia of 9-12 week old mice using PBS (without Mg^2+^ and Ca^2+^) supplemented with 5 uM EDTA and 1% fetal calf serum. Immunostained bone marrow cells were analyzed using gating strategies as described previously by Meinders et al 2014. Briefly, bone marrow cells were gated using cell surface markers c-Kit (BD Pharm anti-CD117 PerCP-Cy5, Cat# 560557), mouse linage antibody cocktail (BD pharm APC mouse linage antibody cocktail, Cat# 558074), CD31 (BD Pharm anti-CD31 PE-Cy7, Cat# 561410), CD41 (BD Pharm anti-CD41 PE, Cat# 558040), CD42b (Novus Biological anti-GPIb alpha Antibody-HIP1 FA750, Cat# NB500-511), CD61 (BD Pharm anti-CD61 FITC, Cat# 561911). Flow cytometry analysis was performed on a BD Symphony A5 unit. c-Kit^+^ MK cells were analyzed for CD42b vs CD41 or CD61 vs CD31 to define MK sub-populations I – V, similar to Meinders et al 2014, except that sub-populations I-V were not further separated by cell size indicator SSC-A (high vs low). Cell ploidy was measured by DNA staining with Hoechst 33342 (Thermo Scientific, Cat# 23491-52-3). Differences in ploidy between Myc-WT and Myc+/-MKI-MKV sub-populations were analyzed using the 2-tailed t test at the 95% confidence level.

### γH2AX Immunofluorescence, drug treatments and confocal microscopy of MYC-ER cells

γH2AX Immunofluorescence staining was performed as described previously (Rehman et al, NAR, 2018). Briefly, approximately 1x10^4^ cells of HOMycER12(44) cells were plated for one hour on Ibidi 15 μ-Slide angiogenesis slides (Ibidi-treat, 81506) and then treated with 4-hydroxy-tamoxifen (200 nM, Sigma-Aldrich), etoposide (25 mM, Sigma Aldrich) or camptothecin (10 mM, Sigma-Aldrich) as indicated, followed by fixation with 4% paraformaldehyde for 20 min at room temperature. Anti-phospho-histone H2AX (ser139), mouse monoclonal antibody (clone JBW301, sc-40; Sigma-Aldrich) was used at 1:200 dilution. Anti-mouse IgG secondary antibody labeled with Alexa 488 (Invitrogen) was used for detection (1: 2000). Cells were mounted in DAPI (Sigma-Milipore) and were imaged with Zeiss a LSM880 Multi-Photon Confocal Microscope. Red pixels of unprocessed images were quantified with Photoshop histogram function. Statistics were calculated using Prism 9.

### Analysis of of γH2AX and Myc protein levels in tumors arising in p53-KO mice

The tissue sections were deparaffinized and rehydrated through xylenes and descending gradient alcohol. Heat-mediated antigen retrieval was performed using a pressure chamber (Pascal; Dako, Carpinteria, CA) with pH 6 citrate buffer (DAKO), and endogenous peroxidase activity was blocked by incubation in 3% H_2_O_2_ for 10 min. The slides were then incubated with rabbit monoclonal anti-c-Myc antibody (Abcam, Cambridge, MA; clone Y69) at 1:250 for 60 min at room temperature or rabbit monoclonal anti-gamma H2AX (Abcam; clone EP845(2)Y) at 1:2000 for 30 min at room temperature in a Dako Autostainer. The antigen-antibody reaction was detected with Dako EnVision + Rabbit HRP (Dako), labeled with DAB+ (3, 3ʹ-Diaminobenzidine; Dako) and counterstained with hematoxylin. Negative controls were performed by omitting the primary antibody and rabbit immunoglobulin (IgG).

### Visiopharm quantitation of protein levels in immunostained histologic sections

To apply digital image analysis, the stained slides were digitalized using NanoZoomer XR digital slide scanner (Hamamatsu, Hamamatsu City, Japan) at × 40 objective magnification with a single-focus layer. The digitalized images were analyzed using Visiopharm software v6.9.1 (Visiopharm, Hørsholm, Denmark). After training the system to identify cells by digitally “painting” nuclei (see example in Supplemental Figure 4B), we defined the nucleus size as 6.5 µm, with a sensitivity of 61% in RGB-G mode. Blue-colored (hematoxylin) tumor nuclei were initially defined, and then brown-colored (DAB) nuclei were separated spectrally. For both c-Myc and H2AX, nuclear staining was evaluated. The DAB intensity of staining was categorized as 0, 1+, 2+, and 3+ according to the threshold of >78.5%, 70-78.5%, 53.5-70%, and <53.5%, respectively, in HDAB-DAB mode. The histoscores were calculated by multiplying the intensity score and proportion score (range, 0-300).

## Acknowledgements

We thank Drs. Shyam Sharan and Eric Batchelor for critical reading and comments on the manuscript, Drs. Markku Miettinien and Zhenfeng Wang for assistance with the initial immunochistochemistry experiments, Shannon Skarshaug and Maxim Semin for RNA quantitative analysis, and Jonathan Keller, Jeff Carrell, and Megan Karwan for advice on analysis of bone marrow megakaryocyte populations and flow cytometry.

## Author Contributions

SM, DL, Sv.P, XB and ZA designed the project, SM and DL wrote the paper and all authors contributed to editing. XB, ZA, VZ, and Sv.P designed and performed the experiments. KY and SMH assisted with Myc IHC. St.P, SMH, SM and DL evaluated tumor histology. JMS generated key independent experimental specimens.

## Data and materials availability

All data is available in the manuscript or the supplementary materials.

## SUPPLEMENTAL FIGURE LEGENDS

**Supplementary Table 1.** Frequency of tumor types in *Myc*-Wt and *Myc*+/- *p53KO* mice.

|  |  | *Myc*<sup>+/-</sup>;*p53KO* |  | *Myc*-WT;*p53KO* |  |
| --- | --- | --- | --- | --- | --- |
|  |  | tumor<br>frequency | median age<br>onset (days) | tumor<br>frequency | median age<br>onset (days) |
| *All |  | 91% | 131 | 98% | 91 |
| Hemangiosarcoma |  | 31% | 105 | 41% | 86 |
| **Lymphoma |  | 17% | 153 | 23% | 99 |
| Hemangiosarcoma+lymphoma |  | 17% | 111 | 20% | 81 |
| ***Other |  | 26% | 83 | 15% | 91 |

\*4-7% death w/o tumor, \*\*~80% thymic primary, \*\*\*adenocarcinoma,sarcoma embryonal carcinoma, PNET

**Figure S1.**
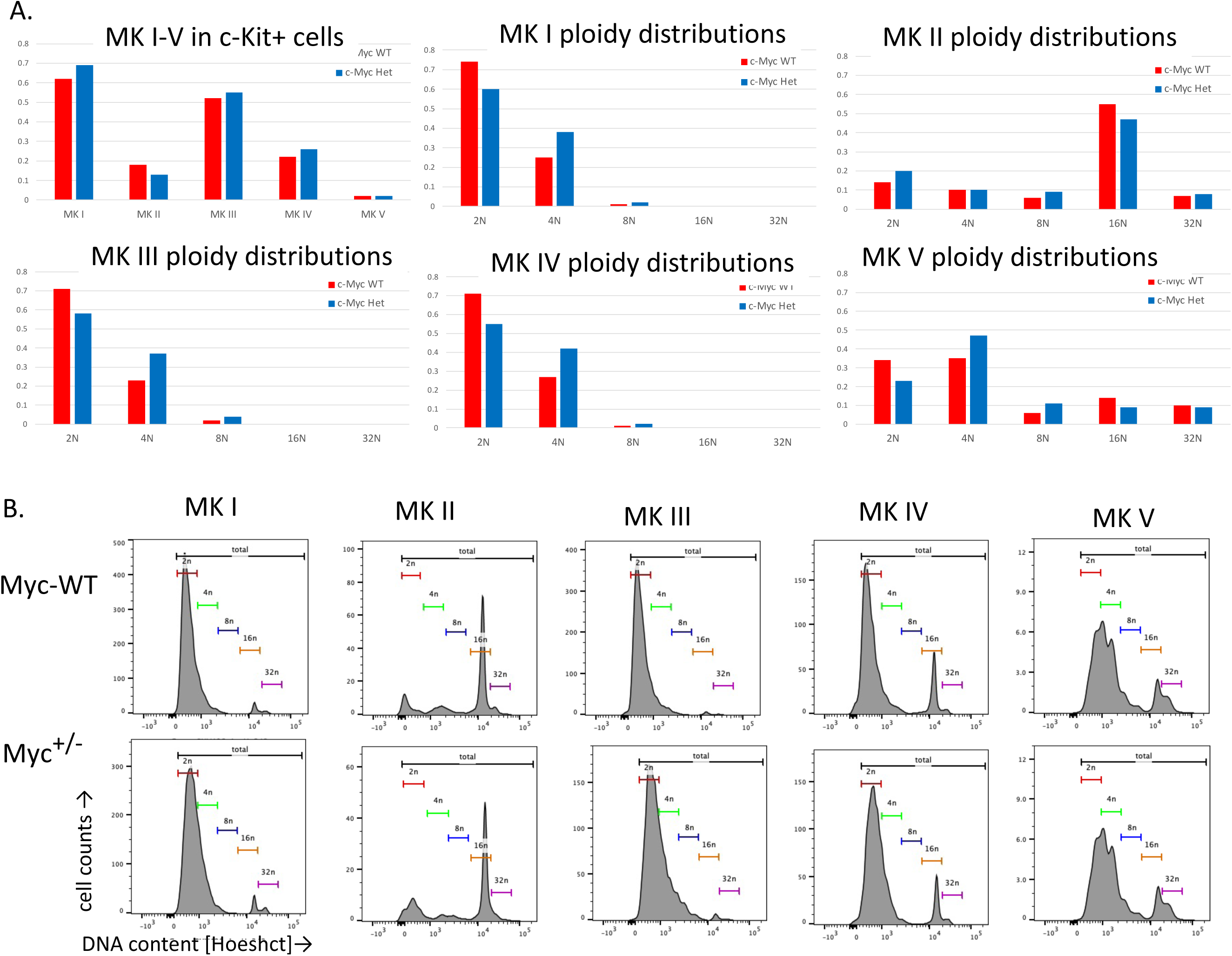
Myc dosage reduction (Myc+/-) does not affect ploidy distribution in a non-neoplastic, normally polyploid population of maturing megakaryocytes. A. Bar graphs comparing distribution of megakaryocyte (MK) subpopulations (MKI-MKV) and the relative DNA content distributions (2N-32N) within a subpopulation for Myc-WT compared to Myc+/-bone marrow cells subjected to flow cytometry (from p53+/-mice; n=4 independent bone marrows from Myc-WT and from Myc+/-mice). No statistically significant differences in either subpopulation distributions or in DNA content profiles within a subpopulation were detected between Myc-WT and Myc+/-megakaryocytes by the 2-tailed t test at the 95% confidence level. B. Representative examples of DNA content profiles from flow cytometry of different megakaryocyte maturation sub-populations MKI-MKV from Myc-WT and Myc+/-mice.

**Figure S2.**
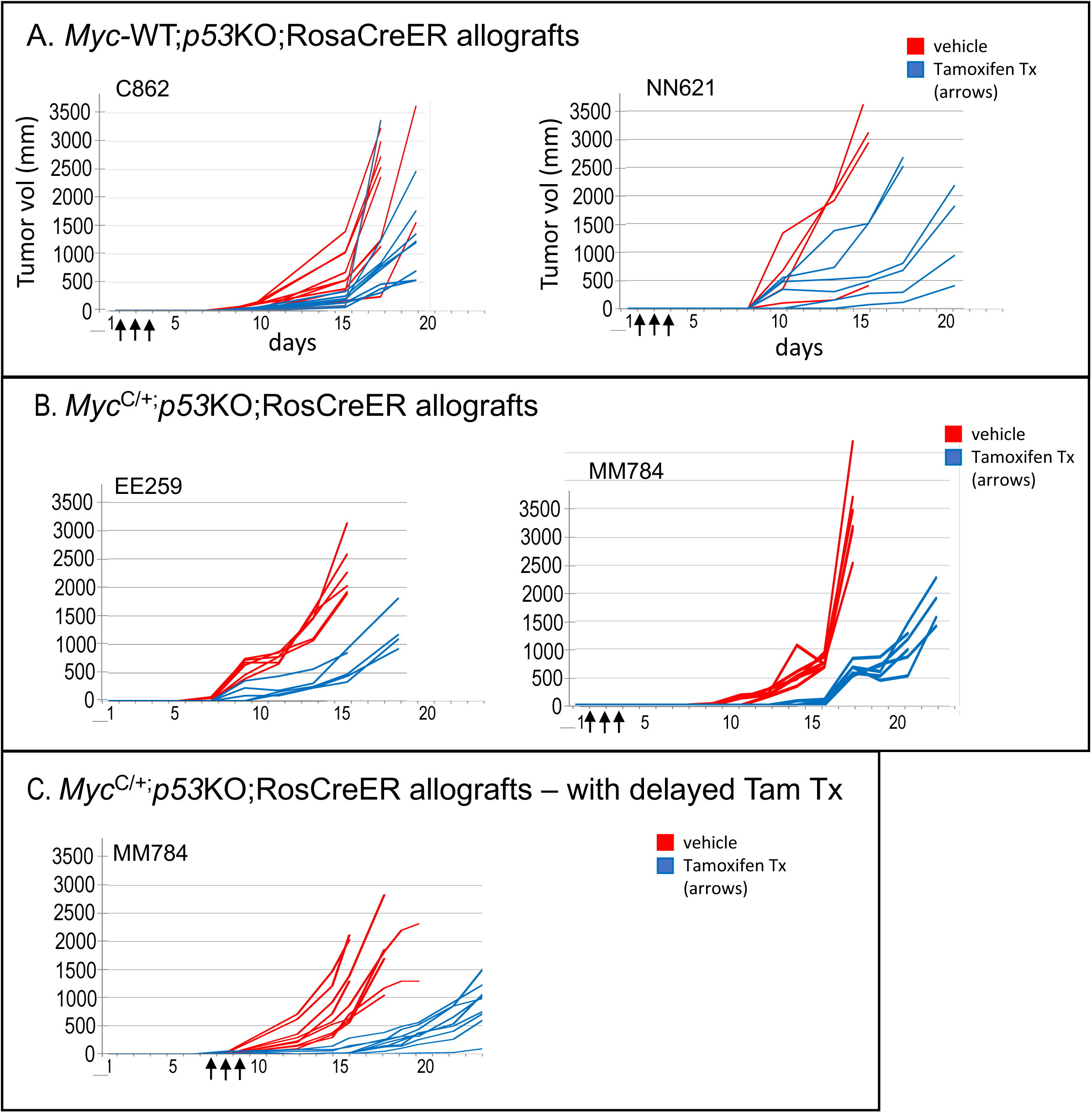
Acute reduction of *Myc* gene dosage in allografted, established *p53KO* thymic lymphomas impairs tumor growth. Primary data for graphs of mean tumor sizes over time, shown in Figure 6. Dissociated cells from *Myc*^+/+^ (C862 and NN621 tumors) or conditional *Myc*^C/+^ (EE259 and MM784 tumors) *p53KO*;RosaCreER+ thymic lymphomas were injected subcutaneously in the flank of nude mice (ranging from 1-5x10^5^ cells) and treated in parallel with either tamoxifen (blue) or vehicle (red) for 3 consecutive days (arrows) beginning at either 1 day (A, B) or 7 days (C) after allografting. Each graph line represents growth plotted for an independent tumor inoculum into flank. A. Tumor allograft growth (mm^3^) for *Myc*-WT C862 and NN621 primary tumors, treated with vehicle or tamoxifen beginning 1 day after allograft. B. Tumor allograft growth (mm^3^) for *Myc*^C/+^ EE259 and MM784 primary tumors, treated with vehicle or tamoxifen beginning 1 day after allograft. C. Tumor allograft growth (mm^3^) for *Myc*^C/+^ MM784 primary tumor, treated with vehicle or tamoxifen beginning after a delay of 7 days after allograft.

**Figure S3.**
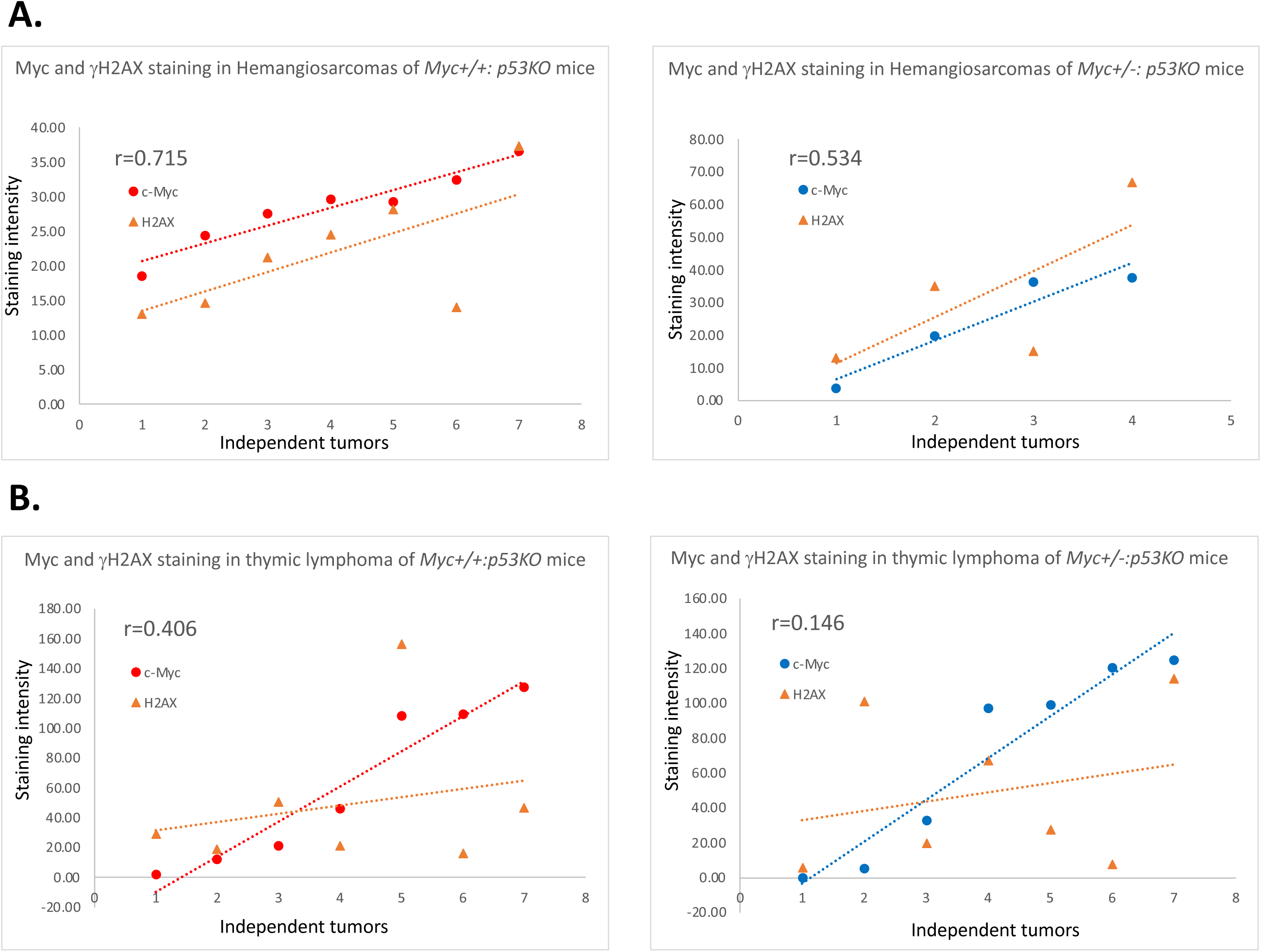
Correlation of Myc and γH2Ax protein levels in hemangiosarcomas but not lymphomas. Correlation of slopes for Myc versus γH2AX in different tumor types for Myc+/- and Myc-WT (as indicated; plotted from data shown in Figure S4). A. Degree of DNA damage as assessed by γH2AX level correlates directly with the Myc level in both Myc+/- and Myc-WT hemangiosarcomas. B. γH2AX/DNA damage and Myc levels correlate poorly in thymic lymphomas. After immunostaining for Myc and H2AX (see Methods), histoscores for both c-Myc and H2AX were generated for each tumor using Visiopharm image analysis (Methods). Tumors are grouped by types (either hemangiosarcoma or thymic lymphoma) and then by c-Myc genotypes (either WT or heterozygous). Pearson’s correlation coefficient (r) of c-Myc and H2AX was calculated within each of the four tumors groups.

**Figure S4.**
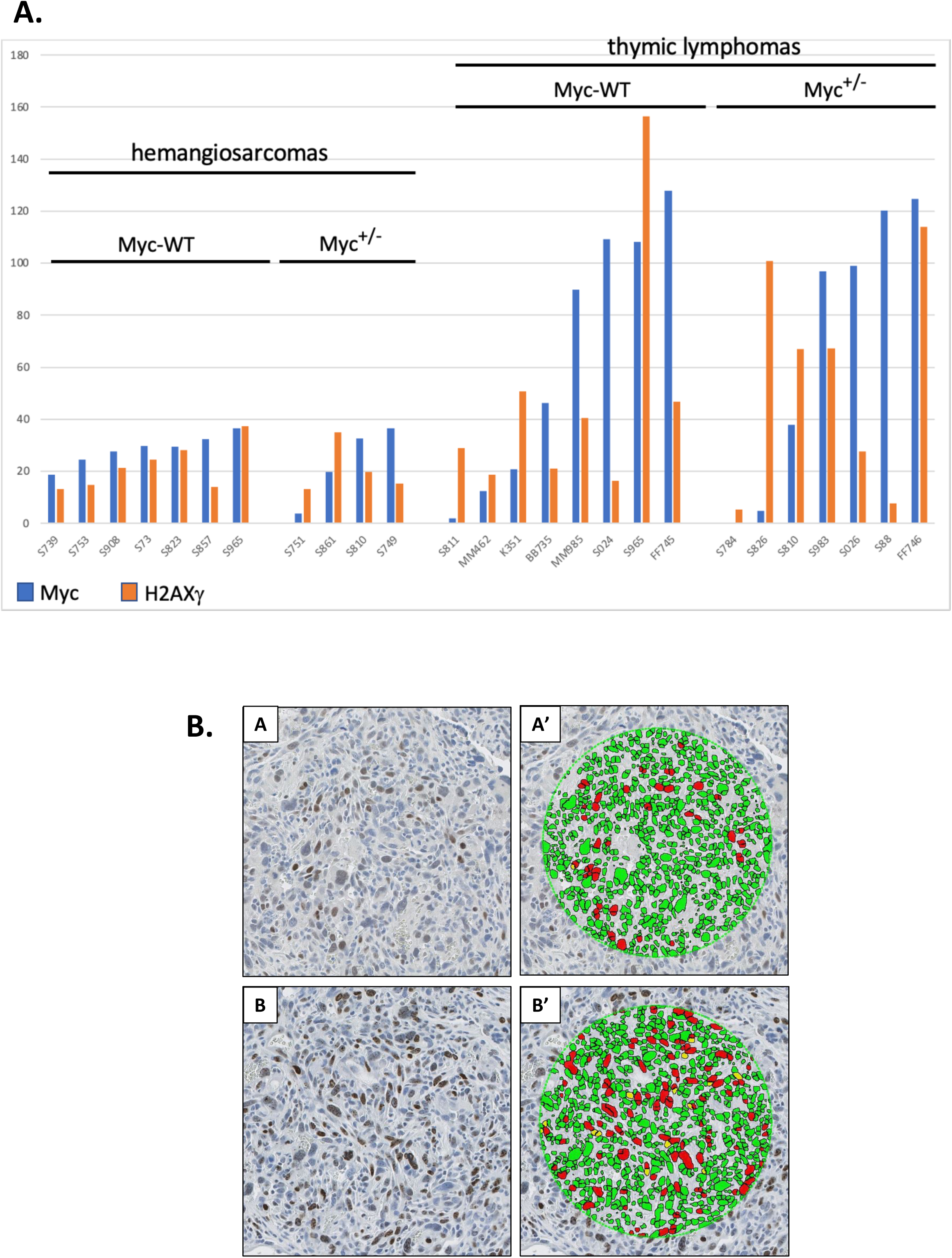
Determination of Myc and γH2Ax protein levels in Myc-WT and Myc+/-hemangiosarcomas and lymphomas from p53KO mice. **A**. Bar graphs of relative Myc and γH2AX protein expression levels determined from immunohistochemical staining using Visopharm software analysis in different p53KO tumors as indicated in graph: for *Myc*-WT and *Myc*^+/-^ p53KO-hemangiosarcomas and for *Myc*-WT and *Myc*^+/-^ thymic lymphomas. **B.** Immunohistochemistry signal quantification using Visiopharm. Examples of “trained” software identification of immunostained cells (A, B) by digitally “painting” nuclei to calculate proportions of nuclear immunostaining (+red nuclei relative to blue in A’, B’). Nuclear size defined as 6.5 µm, with a sensitivity of 61% in RGB-G mode. Blue-colored (hematoxylin) tumor nuclei were initially defined, and then brown-colored (DAB) nuclei were separated spectrally and DAB intensity was categorized as 0, 1+, 2+, and 3+ according to the threshold of >78.5%, 70-78.5%, 53.5-70%, and <53.5%, respectively, in HDAB-DAB mode. See also description in Methods.

**Figure S5.**
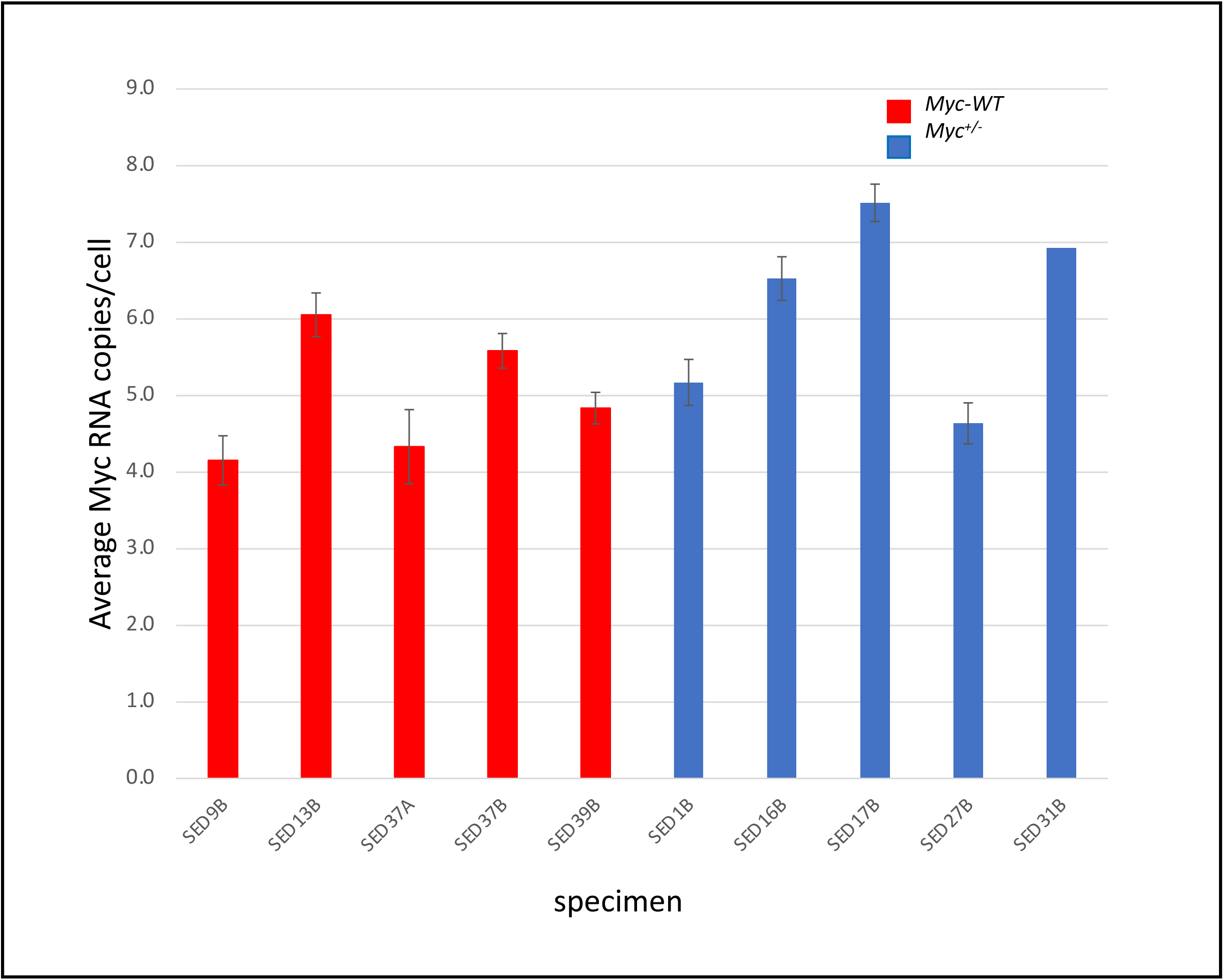
MYC levels must exceed a supraphysiological threshold in spontaneous tumors. *Myc* RNA expression levels are also compensated in spontaneous B-cell lymphomas arising in aged C57BL/6 mice with WT or haploid *Myc* gene dose and wildtype *Trp53* gene. Graph comparing average relative *Myc* RNA (RNA FISH) expression levels and for *Myc*^+/-^ and *Myc*^+/+^ nodal lymphoma cases reported by Hofmann et al. (2015). Bars indicate standard error of the mean (SEM, generated using Visiopharm software).

